# High resolution spatial transcriptome analysis by photo-isolation chemistry

**DOI:** 10.1101/2020.03.20.000984

**Authors:** Mizuki Honda, Shinya Oki, Akihito Harada, Kazumitsu Maehara, Kaori Tanaka, Chikara Meno, Yasuyuki Ohkawa

## Abstract

In multicellular organisms, individual cells are characterized by their gene expression profiles and the spatial interactions among cells enable the elaboration of complex functions. Expression profiling in spatially defined regions is crucial to elucidate cell interactions and functions. Here, we established a transcriptome profiling method coupled with photo-isolation chemistry (PIC) that allows the determination of expression profiles specifically from photo-irradiated regions of whole tissues. PIC uses photo-caged oligodeoxynucleotides for *in situ* reverse transcription. After photo-irradiation of limited areas, gene expression was detected from at least 10 cells in the tissue sections. PIC transcriptome analysis detected genes specifically expressed in small distinct areas of the mouse embryo. Thus, PIC enables transcriptome profiles to be determined from limited regions at a spatial resolution up to the diffraction limit.

## INTRODUCTION

Each cell type in a multicellular organism is characterized by its gene expression profile, which is partly defined by its spatial context. Characterization of whole organs and tissues has recently been advanced by technologies enabling genome-wide expression analysis, as represented by RNA-seq (*1–3*). Properties of individual cells can be studied using single cell RNA-seq by isolating individual cells from dissociated tissues (*4–10*). Expression profiling with respect to spatial information is crucial to determine the characteristics of cells that are controlled, at least in part, by spatial interactions.

Expression patterns of individual or small numbers of genes are classically determined by *in situ* hybridization (ISH) using riboprobes that are visualized by multi-color staining (*11–13*). Localization of hundreds to thousands of transcripts can be analyzed by seqFISH+ and MERFISH by using multiple riboprobe presets coupled with *in situ* sequencing technology (*14, 15*). Unbiased expression profiling associated with spatial information is enabled by spatial transcriptomics and Slide-seq technologies that use multiplexed sequencing with positional barcodes (*16, 17*). Cells located in specific regions of interest (ROIs) can be isolated from specimens by laser microdissection (LMD) and can then be analyzed by RNA-seq (*18*). However, expression profiling of small numbers of cells or single cells from small ROIs remains technically challenging.

By taking advantage of the fact that light is able to change molecular properties with a resolution up to the diffraction limit, we have developed photo-isolation chemistry (PIC), which enables transcriptional profiles of photo-irradiated cells only to be determined. We demonstrate that PIC can provide detailed gene expression profiles for several tens of cells in small ROIs in mouse embryonic tissues. PIC identifies genes uniquely expressed in spatially distinct areas with a resolution up to the diffraction limit.

## RESULTS

### Establishment of PIC expression analysis

To understand multicellular systems within a spatial context, we attempted to develop a gene expression profiling method for small areas with high spatial resolution. The method, termed photo-isolation chemistry (PIC), takes advantage of photo-caged oligodeoxynucleotides (caged ODNs) for the amplification of cDNAs in response to photo-irradiation (Fig. 1a). Experimentally, 1st strand cDNAs are synthesized *in situ* by applying both the caged ODNs and reverse transcriptase onto the tissue sections. Optionally, regions of interest (ROIs) can be precisely defined by immunostaining with antibodies against regional markers. The caged moieties are then cleaved from the ODNs by specific wavelength photo-irradiation under a conventional fluorescence microscope. The whole specimen is then lysed with protease and cDNA:mRNA hybrids extracted. Only the uncaged libraries can be amplified by CEL-seq2 (*19*), a highly sensitive single cell RNA-seq technology. In this manner, only gene expression from photo-irradiated ROIs is detected.

**Fig. 1.**
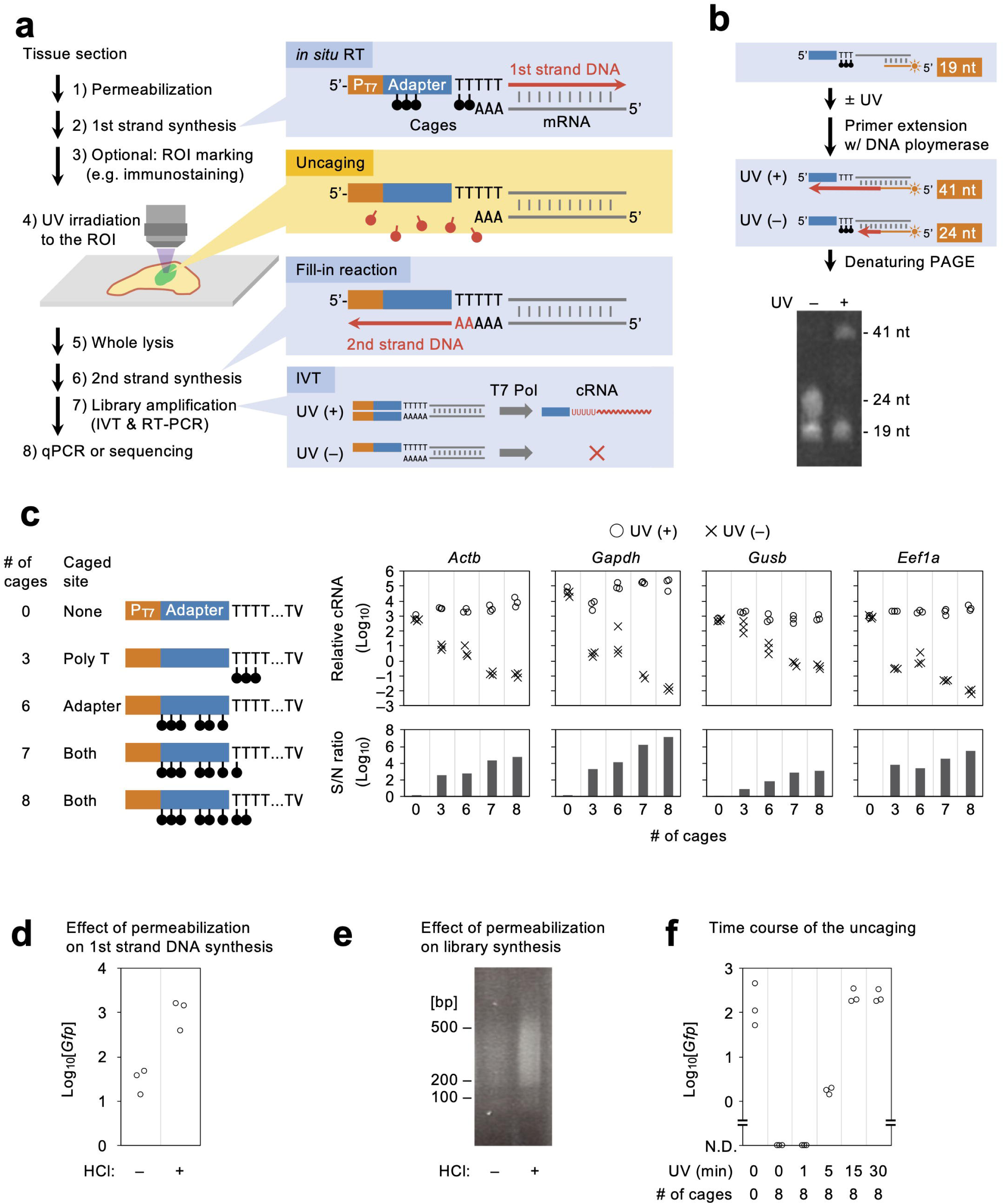
Establishment of the PIC expression profiling method. **a**, Schematic overview of expression profiling with PIC. P_T7_, T7 promoter; T7 Pol, T7 polymerase. **b**, Strategy (top) and the result (bottom) of primer extension experiments to suppress read-through of DNA polymerase I at the NPOM-caged dT sites (black circles) without UV irradiation. **c**, Various caged ODNs were examined for RNA-seq library preparation with (circles) or without (crosses) photo-irradiation before quantifying expression of *Actb, Gapdh, Gusb*, and *Eef1a* genes. **d,e**, Effect of HCl permeabilization before *in situ* RT in GFP-expressing NIH/3T3 cells was evaluated by the yield of *Gfp* cDNA (**d**) and the size (**e**) of sequence libraries. **f**, GFP-expressing NIH/3T3 cells were photo-irradiated for various times after *in situ* RT, and the yield of *Gfp* cDNA quantified.

A critical issue in the development of PIC is the suppression of cDNA amplification from non-irradiated regions. 6-nitropiperonyloxymethyl deoxythymidine (NPOM-dT) was chosen to synthesize the caged ODNs because chain extension by DNA polymerases pauses at NPOM-dT sites in template strands, but resumes upon photo-irradiation of a wavelength around 365 nm (*20*). DNA polymerase I, which is used for 2nd strand DNA synthesis in CEL-seq2, processes through single NPOM-dT sites irrespective of photo-irradiation (*21, 22*). To block DNA polymerase I read-through in PIC, we tried to insert multiple cages in the template strand (Fig. 1b). ODNs harboring triplet NPOM-dTs were annealed with fluorescent dye-labeled primers (19 nt) and chain elongation by DNA polymerase I was examined with or without photo-irradiation. As a result, fully extended complementary strands (41 nt) were produced after photo-irradiation (352–402 nm for 15 min). In contrast, extension was paused at the NPOM-dT triplet (24 nt) site in the absence of photo-irradiation, suggesting that repetitive NPOM-dT insertion effectively suppresses read-through of DNA polymerase I. Based on these results, we designed several caged ODNs and evaluated their signal-to-noise (S/N) ratios by comparing the amounts of amplified libraries with and without photo-irradiation (Fig. 1c). Total RNAs from NIH/3T3 cells (40 ng) were reverse transcribed with caged ODNs and photo-irradiated or not under the fluorescence microscope (352–402 nm for 15 min). The cDNAs were then amplified by 2nd strand synthesis and *in vitro* transcription reactions. Resulting cRNAs of house-keeping genes were quantified by RT-PCR to calculate the S/N ratio. The S/N ratio of non-caged ODNs was nearly equal to 1, indicating that 365 nm UV irradiation was minimally harmful to the integrity of nucleotides. A substantial amount of background was detected by insertion of a NPOM-dTs triplet at the 5′ terminal of the poly-T, with the S/N ratios ranging from 10^0.96^ (*Gusb*) to 10^3.83^ (*Eef1a*). Insertion of six NPOM-dTs in the adaptor sequence elevated the S/N ratios, ranging from 10^1.90^ (*Gusb*) to 10^4.11^ (*Gapdh*). Additional insertion of a single NPOM-dT into the poly-T sequence increased the S/N ratios, ranging from 10^2.89^ (*Eef1a*) to 10^6.27^ (*Actb*). One further NPOM-dT inserted into the poly-T sequence further suppressed the background, with the S/N ratios ranging from 10^3.12^ (*Gusb*) to 10^7.13^ (*Gapdh*). This last caged ODN was the most effective at suppressing background and was therefore used for the experiments described below.

We also examined conditions to enhance the detection sensitivity of PIC. Hydrochloric acid (HCl) is used to enhance efficiency of the *in situ* RT reaction (*23*); therefore, the effect of HCl in the PIC platform was examined. GFP-expressing NIH/3T3 cells grown on coverslips (∼5,000 cells) were fixed and permeabilized in the presence or absence of HCl. *In situ* RT with non-caged ODNs was then performed (Fig. 1d). 1st strand cDNA synthesis was detected without HCl treatment but there was a 38-fold greater yield of cDNA with HCl treatment (qPCR for *Gfp* cDNA; Fig. 1e). HCl treatment also increased the amount of sequence library produced, but the length of cDNAs in the library was not affected (∼200–500 bp; Fig. 1f). The optimal duration of photo-irradiation was also examined. GFP-expressing NIH/3T3 cells on coverslips were fixed and permeabilized with HCl, followed by *in situ* RT with caged ODNs. Coverslips were then irradiated with 340–380 nm light for varying times under the fluorescence microscope and cell lysates then subjected to qPCR for *Gfp* cDNA. *Gfp* cDNA was detected at comparable levels after photo-irradiation for 15 and 30 min. This level was decreased by up to 1% following irradiation for 5 min but no cDNAs were detected after irradiation for 1 min. Taken together, the optimal PIC conditions were as follows: specimens were permeabilized with HCl; caged ODNs with eight of NPOM-caged dTs were used for *in situ* RT; and 340–380 nm photo-irradiation for 15 min was used for uncaging. All the experiments described below were performed using these conditions.

### Validation of PIC for ROI-specific profiling

Suppression of the background from non-irradiated regions is crucial for ROI-specific expression profiling with PIC. To evaluate background levels, mixed-species cultures of human- and mouse-derived cell lines [T-47D (human) and *Gfp*-expressing NIH/3T3 (mouse)] were examined to detect species-specific expression after photo-irradiation (Fig. 2a). T-47D and NIH/3T3 cells were separately aggregated and then mixed and adhered onto coverslips. They were then fixed, permeabilized, and *in situ* RT was performed. Both GFP-negative and positive aggregates were UV-irradiated under the fluorescence microscope. The objective lens was ×40, enabling irradiation of circular areas 750 μm in diameter. After irradiation, cell lysates were analyzed by qPCR for species-specific gene expression. When human T-47D cells were photo-irradiated, human *GAPDH* was detected (*n* = 3/4), but mouse *Gapdh* and *Gfp* were not (*n* = 0/4 each; Fig. 2b, lane 5). In contrast, when *Gfp*-expressing NIH/3T3 cells were photo-irradiated, mouse *Gapdh* and *Gfp* were detected (*n* = 4/4 each), but human *GAPDH* was not (*n* = 0/4; Fig. 2b, lane 6), indicating that background from non-irradiated cells was below the detection limit in mixed culture experiments.

**Fig. 2.**
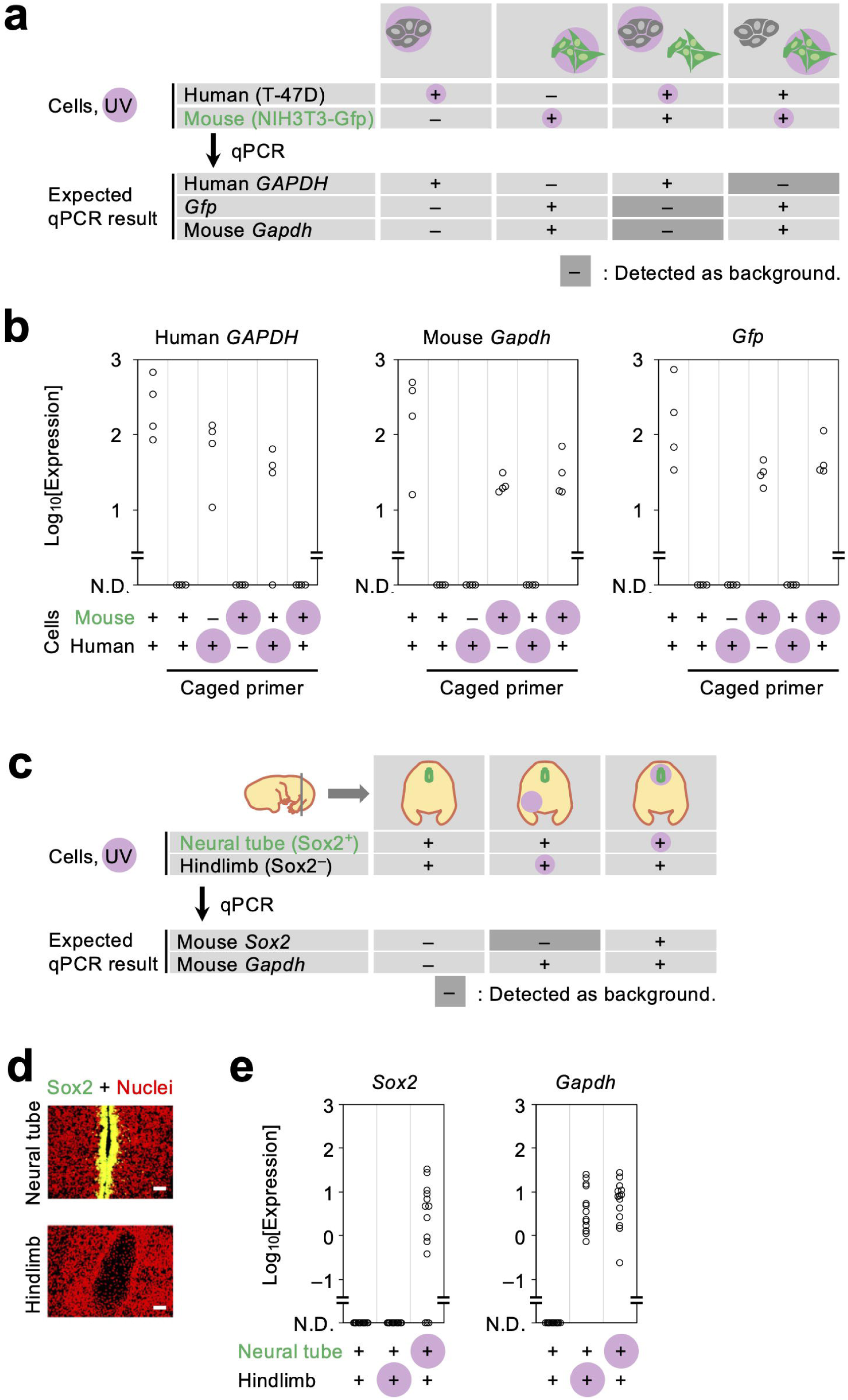
Validation of PIC for ROI-specific profiling. **a**–**e**, Experiments to evaluate the level of background from non-irradiated cells were performed with human-mouse mixed cultures (**a** and **b**) and E14.5 mouse embryos (**c**–**e**). After photo-irradiation (pink) of either human (T-47D, gray) or mouse (GFP-expressing NIH3T3, green) cells (**a**), the cDNAs of human (*GAPDH*) and mouse (*Gapdh* and *Gfp*) genes were examined (**b**). Mouse embryonic sections of Sox2-positive (neural tube, green) or -negative (hindlimb) cells (**c** and **d**) were photo-irradiated and the cDNAs of *Sox2* and *Gapdh* genes were examined (**e**). Scale bars, 50 μm; red in D, nuclei.

We next applied PIC to tissue sections to determine gene expression in photo-irradiated cells (Fig. 2c). Fresh frozen sections of embryonic day (E) 14.5 mouse embryos were fixed, permeabilized, and subjected to *in situ* RT. The sections were then immunostained for SOX2 to label the midline of the neural tube (Fig. 2d). Either SOX2-positive or -negative cells (neural tube or hindlimb, respectively; ×20 lens to irradiate circular areas 1.2 mm in diameter) were then UV irradiated. After irradiation, tissue lysates were collected and analyzed by qPCR for *Sox2* and ubiquitously expressed *Gapdh*. When the neural tube was photo-irradiated, both *Sox2* and *Gapdh* were detected (*n* = 11/14 and 14/14, respectively; Fig. 2e, lane 3). In contrast, when the hindlimb was photo-irradiated, *Gapdh* but not *Sox2* expression was detected (*n* = 14 each; Fig. 2e, lane 2). These data demonstrated that background from non-irradiated cells was not detected either in cultured cells or tissue sections.

### Sensitivity of expression detected by PIC

To examine the detection sensitivity of PIC, varying numbers of cells were photo-irradiated for expression analysis. *Gfp*-expressing NIH/3T3 cells were sparsely inoculated onto coverslips (10,000 cells), followed by fixation, permeabilization, *in situ* RT, and photo-irradiation. The number of irradiated cells was adjusted by changing the magnitude of the objective lens as follows: 2,392–2,766 cells (×2.5 lens), 596–860 cells (×5), 182–239 cells (×10), 53–66 cells (×20), and 17–25 cells (×40). After irradiation, the cell lysates were analyzed by qPCR for *Gfp* and *Gapdh*. The smallest numbers of cells capable of detecting *Gfp* and *Gapdh* were 20 and 17, respectively (Fig. 3a). The quantified expression levels became larger with increasing cell number. cDNAs were not detected in the absence of photo-irradiation (*n* = 0/4).

**Fig. 3.**
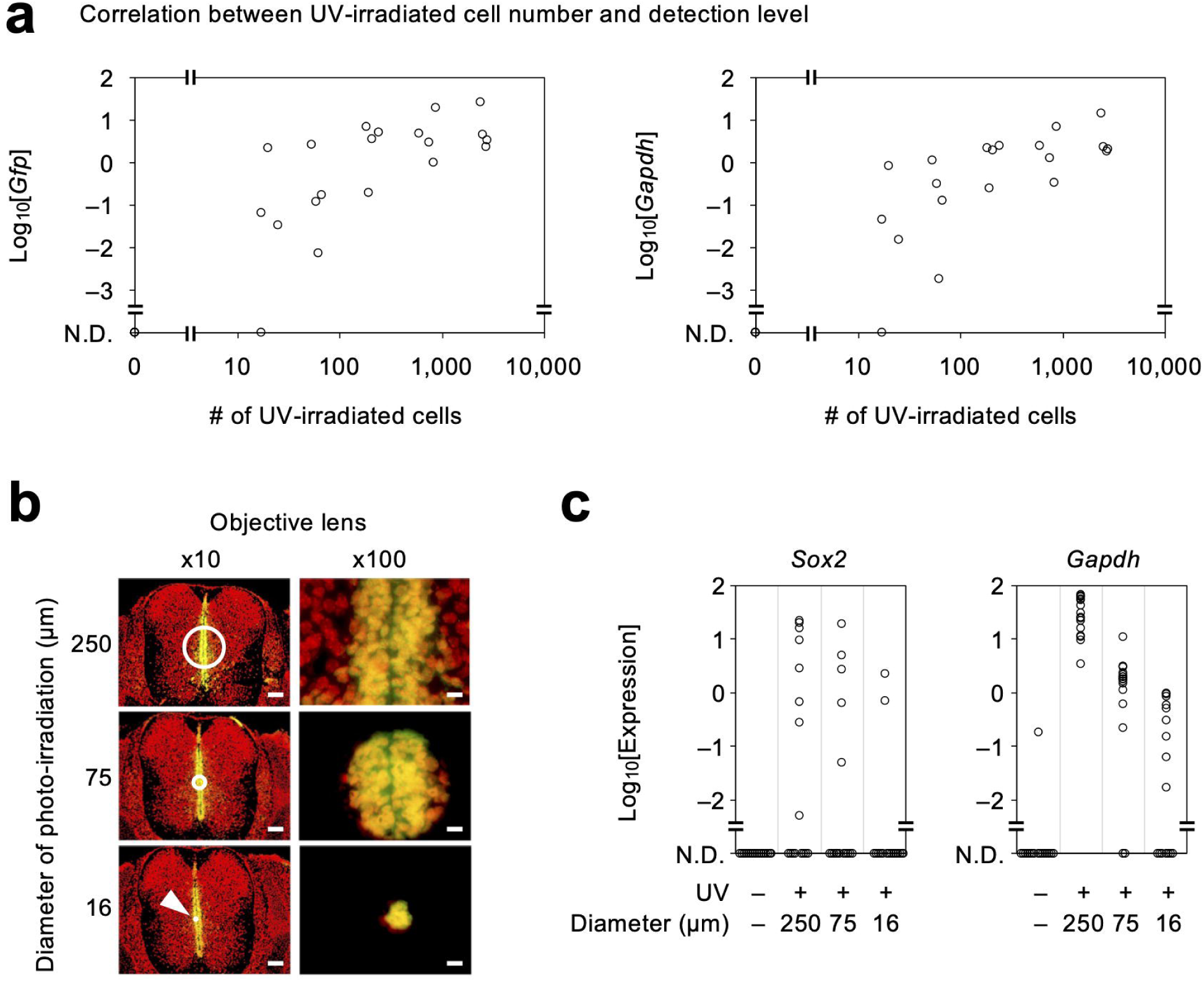
Sensitivity of PIC for low numbers of cells. **a**, Various numbers of GFP-expressing NIH3T3 cells were photo-irradiated and evaluated for levels of *Gfp* and *Gapdh* cDNAs. **b,c**, Areas of various sizes in the E14.5 neural tube (**b**) were photo-irradiated and evaluated for levels of *Sox2* and *Gapdh* cDNAs (**c**). Scale bars in **b**, 100 μm (left) and 10 μm (right).

Detection sensitivity was also evaluated in tissue sections. Frozen E14.5 mouse embryo sections were subjected to *in situ* RT and immunostaining for SOX2. Photo-irradiation of the neural tube was performed with a ×100 objective lens. The sizes of the areas photo-irradiated were adjusted using the fluorescent field diaphragm of the fluorescence microscope, which was changed in a stepwise manner as follows: φ 250 μm (∼ 200 cells), φ 75 μm (∼ 80 cells), or φ 16 μm (∼ 10 cells) (Fig. 3b). After irradiation, the tissue lysates were analyzed by qPCR for *Sox2* and *Gapdh*. A photo-irradiation diameter of at least 16 μm was able to detect *Sox2* and *Gapdh* expression (*n* = 2/16 and 9/16, respectively; Fig. 3c). A photo-irradiation diameter of at least 75 μm was able to detect expression of both genes more frequently (*n* = 5/16 and 14/16, respectively). For a photo-irradiation diameter of 250 μm, *Sox2* was detected in half and *Gapdh* in all samples (*n* = 8/16 and 16/16, respectively). These results indicate that PIC is able to detect gene expression from ten or more cells in tissue sections.

### RNA-seq with PIC

In the developing neural tube, multiple subtypes of interneurons and motor neurons are generated along the dorsoventral axis (*24*). Such neural tube patterning is established by opposing morphogen gradients of BMP/WNT and SHH proteins from dorsal and ventral domains, respectively (*25, 26*). Although a number of genes have been identified to be expressed along the dorsoventral axis by histological methods such as ISH (*27*), genome-wide expression profiling of each domain remains challenging because of the difficulty of isolating such small numbers of cells in a precise manner. We performed PIC to isolate expression profiles from small domains of the neural tube. Following on from our previous experiments (Fig. 3b), E14.5 mouse sections were subjected to *in situ* RT and immunostaining for SOX2. Three distinct domains of the neural tube were independently photo-irradiated (dorsolateral, mediomedial, and ventromedial sites; *n* = 6, 4, and 6, respectively; Fig. 4a) using a ×100 objective lens with a fluorescence field diaphragm (irradiation diameter = 75 μm). As controls, we prepared non-irradiated samples (*n* = 4) and samples in which larger areas were irradiated, centered on the midline of the neural tube (φ 250 and 5,000 μm using ×100 and ×5 lenses, respectively, with an open diaphragm; *n* = 4 each). The specimens were then lysed and libraries amplified and sequenced. On average, sequencing generated approximately ten million reads per sample, irrespective of irradiation size (1.0×10^7^, 5.8×10^6^, and 1.0×10^7^ reads for irradiations of φ 5,000, 250, and 75 μm, respectively; Fig. 4b). In comparison, the number of reads from non-irradiated samples was small (4.3×10^4^ reads), indicating that background reads were rarely included in the reads of photo-irradiated samples. In the photo-irradiated samples, more than half of the reads uniquely mapped to the mouse reference genome (Fig. 4c). The number of gene-assigned reads corrected by unique molecular identifiers (UMIs) was dependent on the irradiation size (4.0×10^5^, 1.9×10^5^, and 4.0×10^4^ reads for φ 5,000, 250, and 75 μm irradiations, respectively; Fig. 4d). Remarkably, more than ten thousand protein-coding genes were detected, even in the smallest areas irradiated (1.6×10^4^, 1.4×10^4^, and 1.0×10^4^ genes for φ 5,000, 250, and 75 μm irradiations, respectively, out of 22,378 Ensembl protein-coding genes; Fig. 4e). We next examined whether the transcriptional profile included spatially specific gene expression. Dimension reduction with uniform manifold approximation and projection (UMAP) revealed that expression profiles from φ 75 μm photo-irradiation areas were clearly separated into three groups according to photo-irradiation site, with the ventromedial and mediomedial genes being closer to each other than dorsolateral genes (Fig. 4f). Differentially expressed genes (DEGs) were detected between dorsolateral versus mediomedial or ventromedial expressed genes (*n* = 198 or 28, respectively) but not between mediomedial versus ventromedial expressed genes (FDR < 0.1; Fig. 4g). The DEGs were largely separated into the following two clusters (Fig. 4h): downregulated in the dorsolateral site (cluster I; *n* = 45) and upregulated in the dorsolateral site, with the latter further branched into two subclusters (clusters II and III; *n* = 61 and 98, respectively). Clusters II and III included several genes known to be expressed in the neural tube, such as *Dcx, Hoxb8, Zic1*, and *Zic4*. ISH experiments demonstrated that the expression levels of these genes were indeed stronger in the dorsal part of the E14.5 neural tube (Fig. 4i), consistent with our expression profiling by PIC.

**Fig. 4.**
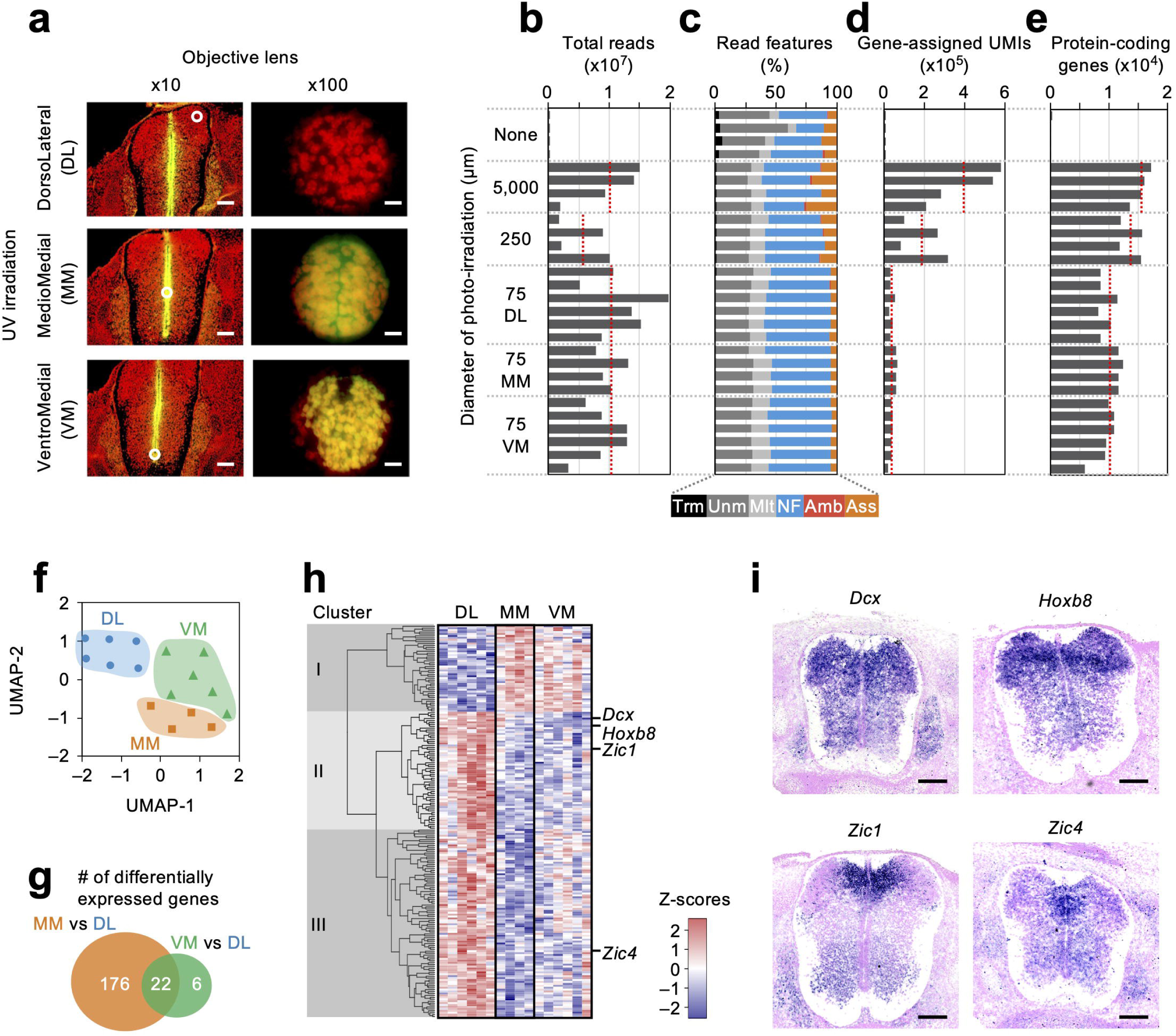
RNA-seq with PIC. **a**, Dorsolateral (DL), mediomedial (MM), or ventromedial (VM) neural tube areas 75 μm in diameter were photo-irradiated and profiled for gene expression with RNA-seq. **b**–**e**, Bar charts showing total read numbers (**b**), read features (**c**), counts of gene-assigned unique molecular identifiers (**d**), and detected protein-coding genes (**e**), with the red dotted lines indicating average values. Abbreviations in **c**: trimmed (Trm), unmapped (Unm), multi-mapped (Mlt), no features (NF), ambiguity (Amb), and gene-assigned (Ass). **f**–**h**, Expression profiles from DL, MM, and VM domains were analyzed by dimension reduction (UMAP) (**f**), and the numbers and clusters of DEGs are shown in a Venn diagram (**g**) and heat maps (**h**), respectively. **i**, Expression patterns of DEGs detected to be stronger in the DL domain by PIC were confirmed by ISH (purple, transcripts; magenta, nuclei). Scale bars in **a**, 100 μm (left) and 10 μm (right); **i**, 200 μm.

## DISCUSSION

We demonstrated that PIC with RNA-seq was able to detect the expression of approximately ten thousand protein coding genes from only tens of cells from the mouse neural tube. Furthermore, ∼200 genes were identified to be expressed with spatial specificity according to the photo-irradiation sites. This technology thus enables gene expression profiles to be obtained from small-scale ROIs. One of the most critical issues of PIC development was to suppress the background amplified from non-irradiated regions. Such background is likely to originate from read-through of DNA polymerase at the NPOM-dT sites of the template strand (*22*). Repetitive insertion of caged nucleotides was effective at decreasing the background to up to less than 0.1% relative to photo-irradiated samples. The use of the optimized caged ODNs reduced the background to under the detection limit of qPCR for *in situ* RT samples from cell cultures and tissue sections. The background of RNA-seq reads was 0.4% relative to φ 75 μm irradiated samples (= an area of 4.4 μm^2^), albeit the area of non-irradiated cells in E14.5 sections was several thousand-fold larger (areas of 10–20 mm^2^). These data indicate that the amount of background is only at a minor level, at least for specimens several millimeters in size. LMD has been used to analyze spatially localized cells by physical isolation of ROIs with a high power laser (*28*). The ROI size and sharpness of the cut edge depends on the laser power and width as well as the stiffness of the specimen. PIC is conceptually different from LMD in the sense that ROIs are photochemically isolated. The spatial resolution of photo-irradiation is up to the diffraction limit; therefore, theoretically, PIC can be applied to submicron-sized ROIs, with the sharpness of the edge being extremely fine. Circular areas can be irradiated using a conventional fluorescence microscope, as shown here. Furthermore, regions with complicated shapes can also be irradiated using pattern illumination systems, such as digital micromirror devices and galvo-based scanning systems (*29, 30*). PIC is thus suitable to analyze cells in complicated fine structures, such as renal glomeruli, which are composed of mesangial cells, podocytes, endothelial cells, Bowman’s capsule, and proximal tubules (*31*). PIC can also be applied to analyze cells in a histologically scattered pattern, such as Sertoli cells in the testes or lymphocytes in inflamed tissues (*32*). Therefore, PIC is able to dissect out detailed characteristics of cells with fine spatial resolution.

To understand cellular characteristics and the spatial interaction of cells at the tissue level, it is necessary to characterize a wide variety of cells by deep expression profiling in combination with high-resolution spatial information. In this context, PIC is able to profile over ten thousand genes from several tens of cells from limited areas. The spatial resolution of PIC is up to the diffraction limit, but the throughput to analyze multiple regions is low. A wide range of cell types can be analyzed by multiplexed profiling with spatial transcriptomics and Slide-seq technologies (*16, 17*). Resulting cell types of interest may be further analyzed by PIC to obtain more detailed expression profiles. Transcriptome in vivo analysis (TIVA) is an alternative method to PIC if targeting living cells (*33*). PIC is also able to detect spatially heterogeneous cells in tissues that appear to be homogenous, such as different spatial domains of the neural tube, as demonstrated in this study. The spatial heterogeneity of transcripts may be analyzed more finely by seqFISH+ at single-molecule resolution. Therefore, PIC is a powerful tool when combined with other spatial transcriptome analyses for understanding multicellular systems.

In the multicellular systems, spatially specific gene expression is orchestrated by chromatin conformation and epigenetic modifications. These regulatory systems have been studied at the genome-wide level to investigate open chromatin, protein-genome binding, and genome methylation, but it remains challenging to perform such analyses on cells located in a limited area. Given that some technologies, such as ATAC-seq, ChIL-seq and PBAT (post-bisulfite adapter tagging), commonly use ODNs as a starting material (*34–37*), caged ODNs provide the advantage of being able to amplify sequence libraries only from photo-irradiated ROIs. Thus, PIC will be able to determine epigenetic landscapes as well as expression profiles in an ROI-specific manner. Multiple cell types that can be juxtaposed or located at a distance from one another can mutually interact. Simultaneous analysis of such interacting cell types will be enabled by the combined use of ODNs caged with NPOM and other caging groups having distinct wavelength-selectivity (*38–40*). After the *in situ* RT with such mixed ODNs, multi-color irradiation and barcode sequencing will separate the sample information. Multi-color PIC will also be useful to compare transcripts showing distinct intracellular localization. Further improvement in PIC sensitivity would be necessary for such innovations.

## ACKNOWLEDGMENTS

We are grateful to the staff of the Ohkawa/Meno Laboratory and we also thank the Advanced Computational Scientific Program of the Research Institute for Information Technology, Kyushu University. We thank Jeremy Allen, PhD, from Edanz Group (www.edanzediting.com/ac) for editing a draft of this manuscript. This work was supported by MEXT/JSPS KAKENHI (19K22398 to M.H., 19H03424 to S.O., JP25116010, JP17H03608, JP17K19356, JP18H04802 and JP18H05527 to Y.O., JP16H01219, JP15K18457 and JP18K19432 to A.H., JP16K18479, JP16H01577, JP16H01550 and JP18H04904 to K.M.), JST PRESTO (JPMJPR1942 to S.O.), and JST CREST (JPMJCR16G1 to Y.O.).

## AUTHOR CONTRIBUTIONS

S.O., and Y.O. conceived PIC and supervised all aspects of the work. M.H. and S.O. performed the experiments. A.H. and C.M. assisted with experiments. K.M. and K.T. analyzed the sequencing data. M.H., S.O., and Y.O. wrote the manuscript. All authors read and approved the final manuscript.

## METHODS

### Cells

NIH/3T3-GFP and T-47D cells were cultured at 37°C in a humidified 5% CO_2_ atmosphere, in high glucose DMEM (Gibco; 11965092) supplemented with 10% fetal calf serum, 1% penicillin/streptomycin (Nacalai Tesque), and 1% L-Glutamin (Nacalai Tesque). To induce the expression of GFP, 1 μg/μl doxycycline (Sigma-Aldrich) was added to NIH/3T3-GFP cells. For sample preparation, cells were dissociated by trypsinization and then inoculated onto gelatinized coverslips either directly or after the formation of cell aggregates by the hanging drop culture method (1,000 cells per 20 μl medium drop for 2–3 days). Total RNA was isolated using an RNeasy Micro Kit (Qiagen).

### Mice

The study was approved by the Animal Care and Use Committee of Kyushu University. Pregnant ICR mice were purchased from Charles River Laboratories Japan and embryos were collected at E14.5. The embryos were immediately embedded in OTC compound (Sakura), and immersed in isopentane/dry ice for flash-freezing. Cryosections at a thickness of 10 μm were mounted on MAS-coated glass slides (Matsunami) and air-dried.

### Caged ODNs

NPOM-caged dT-CE phosphoramidite was purchased from Glen Research (10-1534-95) and used to synthesize caged ODNs with OPC-grade purification by Nihon Gene Research Laboratory.

### PIC protocol

#### Fixation and permeabilization

Cells on coverslips or tissue sections were washed twice with PBS-diethylpyrocarbonate (DEPC), and fixed with 3.7% formaldehyde solution in PBS-DEPC for 10 min at room temperature. Specimens were permeabilized with 5% TritonX-100 in PBS-DEPC for 3 min and then with 0.1 N HCl for 5 min, followed by neutralization with 1 M Tris-HCl pH 8.0 for 10 min at room temperature.

#### In situ RT

Permeabilized specimens were incubated in PBS-DEPC for 5 min at 65°C, and quickly cooled in ice-cold PBS-DEPC. Primer mix [0.5 μl 500 ng/μl caged ODNs, 0.5 μl 10 mM dNTPs (NEB), and 5 μl H_2_O] was also heated to 65°C and quickly cooled to 4°C, before combining with 1st strand mix (2 μl 5× First Strand Buffer, 1 μl 0.1 M DTT, 0.5 μl 40 U/μl RNase Out, and 0.5 μl 200 U/μl Superscript II reverse Transcriptase, all from Invitrogen). The RT reaction mix was applied to the specimens, which were then coverslipped, incubated at 42°C for 60 min, and then heat-inactivated in 70°C PBS for 10 min.

#### Immunostaining

Specimens were blocked with blocking solution [50% Blocking OneP (Nacalai Tesque) in TBST (Tris-buffered saline with 0.1% Tween)] for 10 min at room temperature, incubated with an anti-SOX2 rabbit monoclonal antibody (Cell signaling #23064; 1:1000) overnight at 4°C, then with an Alexa488-conjugated anti-rabbit IgG antibody (Invitrogen; 1:1000; for 1 hr at room temperature). After nuclear staining with TOPRO3 (1:1000 in TBST), specimens were mounted and coverslipped with 90% glycerol containing 1/1000 TOPRO3 and 223 mM 1,4-diazabicyclo[2.2.2.]octane (DABCO).

#### Photo-irradiation

Photo-irradiation of cell cultures and tissue sections for uncaging was performed under a Leica DM5000 B fluorescence microscope illuminated with an EL6000 100 W Hg lamp through a Leica HCX objective lens [2.5×/0.07 Plan (506304), 5×/0.15 PL S-APO (506288), 10×/0.30 PL S-APO (506289), 20×/0.50 PL S-APO (506290), 40×/0.75 PL S-APO (506291), or 100×/1.40-0.70 OIL PL APO (506210)] and a Leica A filter cube (11513873) at a wavelength of 340–380 nm for 15 min unless otherwise indicated. Fluorescence field diaphragms were set at levels 1 and 2 for photo-irradiation of φ16 and 75-μm areas, respectively, with a ×100 lens. Photo-irradiation of solutions in test tubes was performed under a Nikon C1 fluorescence microscope illuminated with a C-HGFI Hg lamp through a 20×/0.45 S Plan Fluor (MRH48230) objective lens and a Semrock DAPI-5060C-NTE filter cube at a wavelength of 352–402 nm for 15 min.

#### Cell lysis

Lysis solution (0.1% Tween and 400 μg/ml Proteinase K in PBS) was loaded onto specimens and incubated for 30 min at 55°C. cDNA:mRNA hybrids were purified with a Qiagen MinElute PCR Purification kit and eluted with H_2_O.

#### 2nd strand DNA synthesis

The eluted cDNA:mRNA hybrids (15 μl) were combined with second strand mix [2 μl 5× First Strand Buffer (Invitrogen), 2.31 μl Second Strand Buffer (Invitrogen), 0.23μl 10 mM dNTPs (NEB), 0.08 μl 10 U/μl *E. coli* DNA ligase (Invitrogen), 0.3 μl 10 U/μl *E. coli* DNA polymerase (Invitrogen), and 0.08 μl 2U/μl *E. coli* RNaseH (Invitrogen)], and incubated for 2 hr at 16°C. The double-stranded cDNAs were purified with AMpure XP beads (Beckman Coulter) and eluted with H_2_O.

#### In vitro transcription

The eluted double-stranded cDNAs (6.4 μl) were combined with *in vitro* transcription mix [1.6 μl each A/G/C/UTP solution, 1.6 μl 10× T7 reaction buffer, and 1.6 μl T7 enzyme from the MEGAscript T7 Transcription Kit (Invitrogen)], and incubated for 17 hr at 37°C. One microliter of 2 U/μl TURBO DNase (Invitrogen) was added and incubated for 15 min at 37°C. Six microliters of ExoSAP-IT PCR Product Cleanup Reagent (Invitrogen) were added and incubated for 15 min at 37°C and then 5.5μl of fragmentation buffer (200mM Tris-acetate pH 8.1, 500 mM KOAc, and 150 mM MgOAc) were added and incubated for 3 min at 94°C. After adding 2.75 μl 0.5 M EDTA pH 8.0, the cRNAs were purified using RNAClean XP beads (Beckman Coulter) and eluted with H_2_O.

#### Library preparation and sequencing

The eluted cRNAs (5 μl) were combined with RT primer mix [1 μl 250 ng/μl randomhexRT primer and 0.5 μl 10 mM dNTPs (NEB)], incubated for 5 min at 65°C, and quickly cooled to 4°C. RT reaction mix was then added [2 μl 5× First Strand Buffer, 1 μl 0.1 M DTT, 0.5 μl 40 U/μl RNase Out, and 0.5 μl 200 U/μl Superscript II reverse Transcriptase (all from Invitrogen)], and incubated for 10 min at 25°C, and further incubated for 1 hr at 42°C. The RT products (9 μl) were combined with PCR mix [1.8 μl each 10 μM RNA PCR primer 1 and 2, 22.5 μl Phusion High-Fidelity PCR Master Mix, and 9.9 μl water], and amplified by PCR (98°C for 30 sec, followed by 11 cycles of 98°C for 10 sec, 60°C for 30 sec and 72°C for 30 sec, with a final extension at 72°C for 10 min). Fragments of 250–500 bp were then purified and size-selected with AMpure XP beads. The quality of the resulting cDNA library was assessed using high sensitivity DNA chips on a Bioanalyzer 2100 (Agilent Technologies) and quantified using a Library Quantification Kit (Clontech), before sequencing on the Illumina HiSeq 1500 platform.

#### Data analysis

The sequence reads were aligned to the GRCm38 reference genome with HISAT2 and analyzed using R packages for read features (featureCount), UMI counting (UMI-tools), dimension reduction (UMAP), DEG analysis (DESeq2), and clustering (heatmap3).

### UV-dependent primer extension

50 μM each of caged ODNs and tetramethylrhodamine (TAMRA)-labeled primer were mixed and incubated for 1 min at 95°C and gradually cool to 25°C during 45 min using a thermal cycler. Half of the annealed oligonucleotides were photo-irradiated for 15 min with 352–402 nm wavelength light as mentioned above. The annealed ODNs with or without photo-irradiation were subjected to the chain extension reaction [final concentration of 1.42 μM annealed ODNs, 1× Second buffer, 0.5 mM dNTPs, and 10 U/μl *E. coli* DNA polymerase I (Invitrogen)] for 2 hr at 16°C, followed by heat denaturation for 1 min at 98°C. Reaction aliquots of 5 μl were electrophoresed on denaturing urea polyacrylamide gels (15%), and TAMRA fluorescence was detected by a UV transilluminator (TOYOBO, FAS-201).

### qPCR

Quantification of several genes in sequence libraries was performed by real-time PCR using NEBnext PCR Master Mix (NEB) supplemented with SYBR Green (Applied Biosystems). Other qPCR experiments were performed using TaqMan Gene Expression Master Mix (Applied Biosystems).

### ISH

Digoxigenin-labeled riboprobes were synthesized with DIG RNA Labeling Mix (ROCHE), Thermo T7 RNA Polymerase (ToYoBo), and gene templates cloned using T7 promoter-containing PCR primer sets and mouse embryonic brain cDNA pools. Fresh-frozen sections of E14.5 mouse embryos were washed twice with PBS-DEPC, and fixed with 4% paraformaldehyde for 12 hr at 4°C. The sections were then serially treated with 6% H_2_O_2_ for 20 min, 10 μg/ml proteinase K solution for 5 min, and post-fix solution (4% paraformaldehyde, 0.2% glutaraldehyde, and 0.1% Tween in PBS-DEPC) for 20 min at room temperature. After incubation with prehybridization mix (50% formaldehyde, 5× SSC pH 5, 150 μg/ml yeast RNA, 150 μg/ml heparin, 1% SDS, and 0.1% Tween) for 30 min at 65°C, hybridization was performed overnight with the same solution containing a digoxigenin-labeled riboprobe. Sections were then washed several times with SSC of increasing stringency and then incubated with alkaline phosphatase-conjugated anti-digoxigenin antibodies (Roche). Nitro blue tetrazolium and 5-bromo-4-chloro-3-indolyl-phosphate (Roche) were used for the colorimetric detection of alkaline phosphatase activity, followed by nuclear staining with 4’,6-diamidino-2-phenylindole (DAPI, Invitrogen). Both bright field and nuclear staining images were separately taken using a Keyence BZ-X700 All-in-One microscope.

## DATA AVAILABILITY

Deep-sequencing data in this study have been deposited in the Gene Expression Omnibus (GEO) under the accession code GSE143413.

## REFERENCES

1. D. Alpern et al., BRB-seq: ultra-affordable high-throughput transcriptomics enabled by bulk RNA barcoding and sequencing. Genome biology 20, 71 (2019).

2. J. Z. Levin et al., Comprehensive comparative analysis of strand-specific RNA sequencing methods. Nature methods 7, 709–715 (2010).

3. D. Parkhomchuk et al., Transcriptome analysis by strand-specific sequencing of complementary DNA. Nucleic acids research 37, e123 (2009).

4. T. Hashimshony, F. Wagner, N. Sher, I. Yanai, CEL-Seq: single-cell RNA-Seq by multiplexed linear amplification. Cell reports 2, 666–673 (2012).

5. T. Hayashi et al., Single-cell full-length total RNA sequencing uncovers dynamics of recursive splicing and enhancer RNAs. Nature communications 9, 619 (2018).

6. S. Islam et al., Highly multiplexed and strand-specific single-cell RNA 5’ end sequencing. Nature protocols 7, 813–828 (2012).

7. S. Picelli et al., Full-length RNA-seq from single cells using Smart-seq2. Nature protocols 9, 171–181 (2014).

8. A. B. Rosenberg et al., Single-cell profiling of the developing mouse brain and spinal cord with split-pool barcoding. Science 360, 176–182 (2018).

9. M. Saikia et al., Simultaneous multiplexed amplicon sequencing and transcriptome profiling in single cells. Nature methods 16, 59–62 (2019).

10. Y. Sasagawa et al., Quartz-Seq: a highly reproducible and sensitive single-cell RNA sequencing method, reveals non-genetic gene-expression heterogeneity. Genome biology 14, R31 (2013).

11. A. M. Femino, F. S. Fay, K. Fogarty, R. H. Singer, Visualization of single RNA transcripts in situ. Science 280, 585–590 (1998).

12. J. H. Lee et al., Highly multiplexed subcellular RNA sequencing in situ. Science 343, 1360–1363 (2014).

13. S. Shah, E. Lubeck, W. Zhou, L. Cai, In Situ Transcription Profiling of Single Cells Reveals Spatial Organization of Cells in the Mouse Hippocampus. Neuron 92, 342–357 (2016).

14. K. H. Chen, A. N. Boettiger, J. R. Moffitt, S. Wang, X. Zhuang, RNA imaging. Spatially resolved, highly multiplexed RNA profiling in single cells. Science 348, aaa6090 (2015).

15. C. L. Eng et al., Transcriptome-scale super-resolved imaging in tissues by RNA seqFISH. Nature 568, 235–239 (2019).

16. S. G. Rodriques et al., Slide-seq: A scalable technology for measuring genome-wide expression at high spatial resolution. Science 363, 1463–1467 (2019).

17. P. L. Stahl et al., Visualization and analysis of gene expression in tissue sections by spatial transcriptomics. Science 353, 78–82 (2016).

18. S. Nichterwitz et al., Laser capture microscopy coupled with Smart-seq2 for precise spatial transcriptomic profiling. Nature communications 7, 12139 (2016).

19. T. Hashimshony et al., CEL-Seq2: sensitive highly-multiplexed single-cell RNA-Seq. Genome biology 17, 77 (2016).

20. A. Prokup, J. Hemphill, A. Deiters, DNA computation: a photochemically controlled AND gate. Journal of the American Chemical Society 134, 3810–3815 (2012).

21. A. Kuzuya, F. Okada, M. Komiyama, Precise site-selective termination of DNA replication by caging the 3-position of thymidine and its application to polymerase chain reaction. Bioconjugate chemistry 20, 1924–1929 (2009).

22. D. D. Young, H. Lusic, M. O. Lively, A. Deiters, Restriction enzyme-free mutagenesis via the light regulation of DNA polymerization. Nucleic acids research 37, e58 (2009).

23. J. H. Lee et al., Fluorescent in situ sequencing (FISSEQ) of RNA for gene expression profiling in intact cells and tissues. Nature protocols 10, 442–458 (2015).

24. J. Briscoe, S. Small, Morphogen rules: design principles of gradient-mediated embryo patterning. Development 142, 3996–4009 (2015).

25. E. Dessaud, A. P. McMahon, J. Briscoe, Pattern formation in the vertebrate neural tube: a sonic hedgehog morphogen-regulated transcriptional network. Development 135, 2489–2503 (2008).

26. I. Patten, M. Placzek, Opponent activities of Shh and BMP signaling during floor plate induction in vivo. Current biology : CB 12, 47–52 (2002).

27. L. Wine-Lee et al., Signaling through BMP type 1 receptors is required for development of interneuron cell types in the dorsal spinal cord. Development 131, 5393–5403 (2004).

28. M. R. Emmert-Buck et al., Laser capture microdissection. Science 274, 998–1001 (1996).

29. B. W. Avants, D. B. Murphy, J. A. Dapello, J. T. Robinson, NeuroPG: open source software for optical pattern generation and data acquisition. Frontiers in neuroengineering 8, 1 (2015).

30. R. Schuck et al., Multiphoton minimal inertia scanning for fast acquisition of neural activity signals. Journal of neural engineering 15, 025003 (2018).

31. A. R. Kitching, H. L. Hutton, The Players: Cells Involved in Glomerular Disease. Clinical journal of the American Society of Nephrology : CJASN 11, 1664–1674 (2016).

32. R. E. Emerson, T. M. Ulbright, Morphological approach to tumours of the testis and paratestis. Journal of clinical pathology 60, 866–880 (2007).

33. D. Lovatt et al., Transcriptome in vivo analysis (TIVA) of spatially defined single cells in live tissue. Nature methods 11, 190–196 (2014).

34. J. D. Buenrostro, P. G. Giresi, L. C. Zaba, H. Y. Chang, W. J. Greenleaf, Transposition of native chromatin for fast and sensitive epigenomic profiling of open chromatin, DNA-binding proteins and nucleosome position. Nature methods 10, 1213–1218 (2013).

35. A. Harada et al., A chromatin integration labelling method enables epigenomic profiling with lower input. Nature cell biology 21, 287–296 (2019).

36. F. Miura, Y. Enomoto, R. Dairiki, T. Ito, Amplification-free whole-genome bisulfite sequencing by post-bisulfite adaptor tagging. Nucleic acids research 40, e136 (2012).

37. S. A. Smallwood et al., Single-cell genome-wide bisulfite sequencing for assessing epigenetic heterogeneity. Nature methods 11, 817–820 (2014).

38. C. Menge, A. Heckel, Coumarin-caged dG for improved wavelength-selective uncaging of DNA. Organic letters 13, 4620–4623 (2011).

39. A. Rodrigues-Correia, X. M. Weyel, A. Heckel, Four levels of wavelength-selective uncaging for oligonucleotides. Organic letters 15, 5500–5503 (2013).

40. X. Tang et al., Caged nucleotides/nucleosides and their photochemical biology. Organic & biomolecular chemistry 11, 7814–7824 (2013).

